# Single cell RNA-seq reveals profound transcriptional similarity between Barrett’s esophagus and esophageal glands

**DOI:** 10.1101/313049

**Authors:** Richard Peter Owen, Michael Joseph White, David Tyler Severson, Barbara Braden, Adam Bailey, Robert Goldin, Lai Mun Wang, Nicholas David Maynard, Angie Green, Paolo Piazza, David Buck, Mark Ross Middleton, Chris Paul Ponting, Benjamin Schuster-Böckler, Xin Lu

## Abstract

Barrett’s esophagus is a precursor of esophageal adenocarcinoma. In this common condition, squamous epithelium in the esophagus is replaced by columnar epithelium in response to acid reflux. Barrett’s esophagus is highly heterogeneous and its relationships to normal tissues are unclear. We investigated the cellular complexity of Barrett’s esophagus and the upper gastrointestinal tract using RNA-sequencing of 2895 single cells from multiple biopsies from four patients with Barrett’s esophagus and two patients without esophageal pathology. We found that uncharacterised cell populations in Barrett’s esophagus, marked by *LEFTY1* and *OLFM4*, exhibit a profound transcriptional overlap with a subset of esophageal cells, but not with gastric or duodenal cells. Additionally, SPINK4 and ITLN1 mark cells that precede morphologically identifiable goblet cells in colon and Barrett’s esophagus, potentially aiding the identification of metaplasia. Our findings reveal striking transcriptional relationships between normal tissue populations and cells in a premalignant condition, with implications for clinical practice.

## Introduction

At least 80% of cancers arise from epithelial cells. In many tumours a change in cell type, referred to as metaplasia, is a key step in cancer initiation. Barrett’s esophagus (BE) is an example of metaplasia in the distal esophagus and affects 1 in 50 people^1^. BE is defined as replacement of squamous epithelium by columnar epithelium, and it gives a 30-fold increased risk of developing esophageal adenocarcinoma (EAC) which is highly fatal^2-4^. BE is associated with gastroesophageal reflux disease, suggesting it occurs in response to a chronically inflamed environment^5^. Remarkably, several anatomically distant cell types are also identifiable in BE, most commonly intestinal goblet cells but also Paneth and pancreatic acinar cells, among others^6-8^.

This apparent plasticity in BE has obscured its relationship with normal gastrointestinal tissues, as no normal gastrointestinal tissue has the extent of cellular heterogeneity as BE. The current most widely held view is that BE originates from the stomach^9,10^, and studies looking for similarities (e.g. in gene or protein expression and cellular appearance) between BE and selected normal tissues - including the intestine, gastric pylorus, gastric corpus and gastric cardia – have found some shared attributes^11^. However, to complicate matters, there is evidence suggesting BE may originate directly from native esophageal cells^12-15^, from recruitment of circulating stem cells^16^, or from reactivation of dormant progenitor cells *in situ*^17^. Therefore, BE characterizations that focus on characteristics of gastric or intestinal columnar epithelia are inherently compromised. A more unbiased and systematic approach to BE characterization in humans is required, not least because BE has been observed after gastric resection^18^, which conflicts with described theories of BE development, rodents and humans have differences in their gastrointestinal tract (such as rodents lacking esophageal glands), and similarities between gastric heterotopia and BE.

Single cell RNA-sequencing (RNA-seq), combined with computational methods for functional clustering of cell types, provides an unbiased approach to understanding cellular heterogeneity. Applying this to human tissues is challenging, but is currently the best way to perform unbiased functional assessment of cells from BE and related normal tissues, with the potential to identify unique cell types and propose new markers of specialised cell types in BE, with implications for how BE can be detected and diagnosed.

Given the highly heterogeneous nature of BE, we hypothesised that single cell RNA-seq might clarify the relationships between cells in normal tissues and BE, and indicate whether there are specialised cells in BE with similar functions to cells elsewhere in the non-inflamed gastrointestinal tract. Therefore we applied this approach to biopsies from BE, esophagus, stomach and small intestine (duodenum). This revealed a cell population in BE that expresses the developmental gene (*LEFTY1)* and is distinct from all tested intestinal or gastric cells, but was transcriptionally highly similar to rare columnar cells from normal esophagus.

## Results

### Single cell RNA-seq identifies subpopulations in normal upper gastrointestinal epithelia

To characterise the cell populations in BE, samples were taken from seven patients attending for routine endoscopic surveillance of non-dysplastic BE previously noted to have intestinal metaplasia. From each patient, we took biopsies from BE, adjacent macroscopically normal esophagus (minimum 20mm proximal to BE), stomach (10-20mm distal to the gastroesophageal junction) and duodenum (**Figure 1a**). Individual 2mm biopsies were divided to provide tissue for single cell RNA-seq, bulk tissue RNA-seq and histology in four patients, and bulk tissue RNA-seq and histology in three patients (see **Methods**). Single cells were also prepared from normal esophageal biopsies from two control patients with gastroesophageal reflux disease but no previous or current diagnosis of BE or any other esophageal pathology. All sampled patients were taking regular acid suppression therapy and had no features of esophageal dysplasia or malignancy (**Extended Data Table 1**).

**Figure 1.**
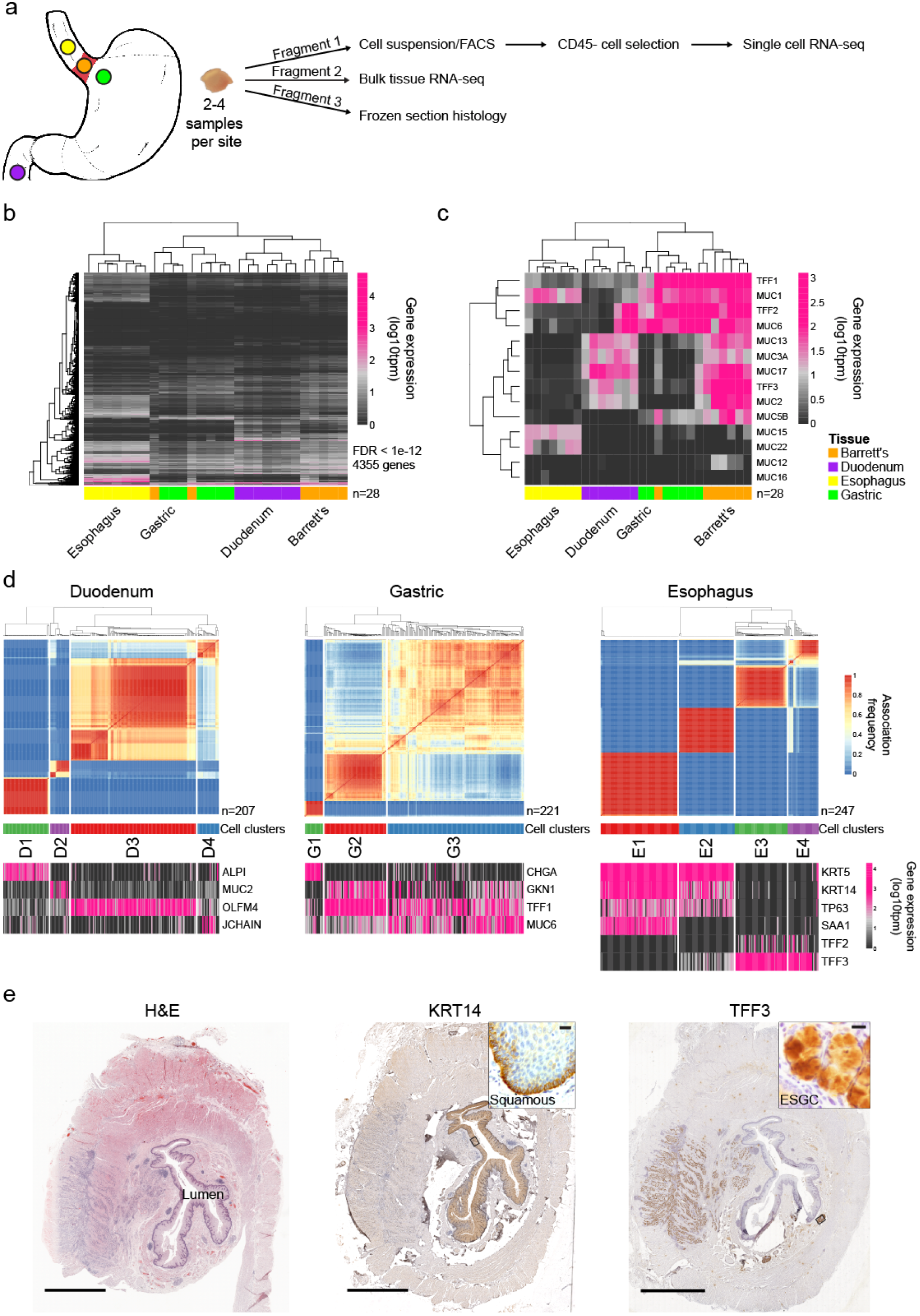
Single cell RNA sequencing identifies cell groups in normal upper gastrointestinal epithelia. **(a)** Endoscopic sampling sites (yellow, esophagus; green, gastric cardia; purple, duodenum; orange, Barrett’s esophagus) with summary of how tissues from patients were used. 2-4 biopsies were taken at each site. Patients without BE were sampled from the lower esophagus 2cm proximal to the squamous-columnar junction. **(b)** From bulk RNA-seq data derived from samples from seven patients with BE, heatmap of genes differentially expressed between any tissue type (FDR < 1e-12) with tissue hierarchy determined by nearest neighbor. Tissue indicated by colours as in **a**. **(c)** From bulk RNA-seq data, heatmap of expression of mucin and trefoil factor genes with tissue hierarchy determined by nearest neighbor, in samples from seven patients with BE. **(d)** Upper panels show the cluster consensus matrices for single cells from normal tissue sites in four BE patients. Blue-to-red colours denote the frequency with which cells are grouped together in 250 repeat clusterings of simulated technical replicates (see Methods). Cell clusters are indicated by coloured bars below the matrices. In lower panels, heatmaps show expression of known functionally relevant genes that were differentially expressed between cell clusters (>4 fold change, FDR <1e-5). **(e)** Representative image of haematoxylin and eosin staining of esophagus taken from an esophagectomy specimen with normal submucosal glands and squamous epithelium (left). An area of adenocarcinoma is present on the left side of each image. Immunohistochemical staining of KRT14 and TFF3 (centre and right, respectively) in adjacent sections from the same specimen with enlargements of representative positively stained regions. ESGC, esophageal submucosal gland complex; squamous, squamous epithelium. Scale bars indicate 5000 μm, inset images 10 μm.

Bulk RNA-sequencing followed by hierarchical clustering of differentially expressed genes in the duodenal, gastric, esophageal and BE samples from the 7 patients with BE showed a clear distinction between squamous (i.e. normal esophagus) and non-squamous (i.e. gastric, duodenum and BE) epithelia (**Figure 1b**). BE samples from all 7 patients had some similarities to duodenal and gastric samples (**Figure 1b**). When a defined list of genes known to distinguish gastrointestinal epithelia^11^ was used in hierarchical clustering, all BE samples appeared most closely related to gastric tissue, consistent with previous studies^9^ (**Figure 1c**).

To gain insights into the cellular heterogeneity of these tissues, we analysed the transcriptomes of 2176 single cells from four BE patients (620 BE cells; 526 adjacent normal esophagus cells; 678 gastric cells; and 352 duodenum cells) and 719 normal esophageal cells from two control patients. A mean of 1.4×10^5^ reads were mapped per cell and a median of 4484 genes/cell were detected (with at least one read per cell) in cells included in the analysis. Based on data from positive (neural RNA diluted to 10pg) and negative (only spike-in control) samples, cells with fewer than 25119 total mapped reads were excluded, leaving a total of 1778 cells for further analysis (see **Methods**).

First, we clustered the cells from each normal tissue type from the BE patients by gene expression (**Figure 1d**). The eleven clusters (D1-D4, G1-G3 and E1-E4, in duodenum, gastric and esophagus samples, respectively) were then annotated on the basis of genes previously characterized as expressed in specific cell types (complete list in **Supplementary Table 1**). In duodenum, these are: intestinal alkaline phosphatase (*ALPI*) expressing enterocytes (D1); mucin 2 (*MUC2*) expressing goblet cells (D2); olfactomedin 4 (*OLFM4*) expressing crypt cells (D3); and some uncharacterized cells expressing Joining Chain Of Multimeric IgA And IgM (*JCHAIN*) (D4). In gastric, these are: chromogranin (*CHGA*) expressing enteroendocrine cells (G1); gastrokinin (*GKN1*) and trefoil factor 1 (*TFF1*) expressing foveolar cells (G2); and mucin 6 (*MUC6*) and *TFF1* expressing mucus neck cells (G3). Of note, the proton pump gene *ATP4A* and the intrinsic factor gene *GIF* were rarely detectable in gastric cells indicating these are cardiac-type gastric samples (**Extended Data Figure 1**).

Interestingly, four clusters were identified in the esophageal samples. Two of these express expected squamous genes (*KRT5*, *KRT14*, *TP63*; clusters E1 and E2) and two express the columnar genes *TFF2* and *TFF3* (clusters E3 and E4). The two squamous clusters can be distinguished by presence (E1) or absence (E2) of acute phase response (*SAA1*) gene expression, representing squamous cells in different states. The expression of *TFF2* and *TFF3* in E3 and E4 is consistent with these cells being from the columnar epithelium of esophageal gland complexes (ESGCs)^19^, an infrequent structure in normal esophagus. For confirmation of this expression pattern, we examined TFF3 and KRT14 protein expression by immunohistochemistry in five normal squamous esophagus resection specimens. As expected, KRT14 was present in squamous epithelium and TFF3 was detected in an esophagus section with clearly defined ESGCs (**Figure 1e**). These results show that single cell transcriptomic analysis can identify gastrointestinal epithelial cell subpopulations, including rare cell populations from ESGCs that cannot be distinguished by conventional RNA-seq.

### Barrett’s esophagus is enriched for *LEFTY1* expressing cells

To identify genes characteristic of distinct BE cell populations we clustered all the BE cells by gene expression (**Figure 2a,** also see **Supplementary Table 1**). The clusters (B1-B4) can be distinguished by expression of *MUC2* (B1; goblet cells); *LEFTY1* (B2 and B3, approximately 71% of BE cells); and *CHGA* (B4; enteroendocrine cells). *KRT7* is expressed similarly across all 4 clusters, consistent with its clinical utility in BE diagnosis. The different expression patterns of MUC2, LEFTY1 and CHGA can also be consistently seen at the protein level in 41 sections from 19 patients (**Extended Data Table 2**); for example, morphologically identifiable goblet cells are positive for MUC2 but not LEFTY1 or CHGA (**Figure 2b**).

The *LEFTY1* expressing cells (B2 and B3; **Figure 2a**) are divided into a larger low proliferating (*MKI67* (Ki67) negative) cluster (B2) and a smaller high proliferating (*MKI67* positive) cluster (B3). LEFTY1, a secreted protein and transforming growth factor beta (TGF-β) superfamily member, is normally expressed in development, where it has roles in left-right asymmetry determination^20^, but little is known about potential roles in adult tissues and it has not previously been associated with BE. *LEFTY1* expression was rare in duodenal and gastric cells, compared to BE (**Extended Data Figure 2**).

**Figure 2.**
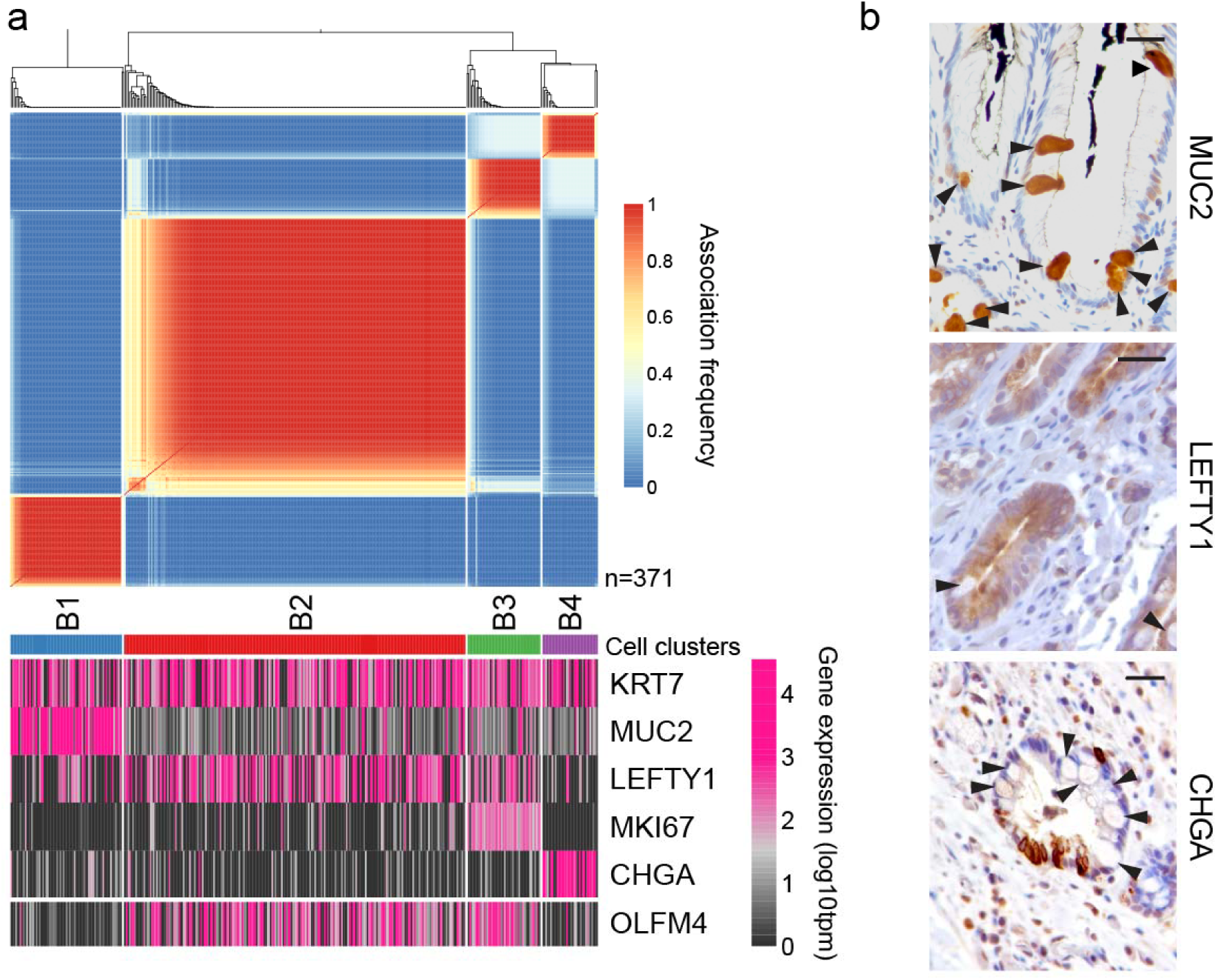
*LEFTY1* and *OLFM4* are mainly expressed in Barrett’s esophagus cells that do not express differentiated secretory cell markers. **(a)** Upper panel, cluster consensus matrix of BE cells from 4 BE patients (n=371). Blue-to-red colours denote the frequency with which cells are grouped together in 250 repeat clusterings of simulated technical replicates (see Methods). Clusters (B1-B4) are indicated by the coloured bars below. Lower panel, heatmaps showing expression of selected functionally relevant genes that are differentially expressed between cell clusters (>4 fold change, FDR <1e-5). **(b)** Immunohistochemical staining of MUC2, LEFTY1 and CHGA in adjacent sections from BE resection specimens. Black arrows indicate goblet cells on all sections (positively stained for MUC2; negative for LEFTY1 and CHGA). Scale bars indicate 50 μm.

### Esophageal gland complexes share a transcriptional profile with Barrett’s esophagus

Taking all cells from BE patients together, the normal tissue cells from the four BE patients separate clearly based on their gene expression, but the BE cells overlap with a sub-set of esophageal cells, as seen in a t-Distributed Stochastic Neighbor Embedding (t-SNE) plot (**Figure 3a**). Clustering by gene expression (by the same method as in Figure 1d) assigned cells to 7 clusters (with brain controls in a separate cluster) (**Figure 3b,c,** also see **Extended Data Figure 3a,b**). Most of these clusters are similar to those identified in the analysis of normal tissue alone (**Figure 1d**) and they can be related to known cell types based on expression of previously characterised genes (**Extended Data Figure 3c,** also see **Supplementary Table 2** for complete list). The majority of duodenal cells fall in the cluster categorised as ‘enterocytes’ (similar to D1), gastric as ‘mucus neck’ (similar to G3), and a substantial proportion of esophageal cells are in the ‘squamous’ cluster (similar to E1/E2) (**Figure 3c**). Some esophageal cells, BE cells and a few duodenal cells fall into a ‘goblet’ cluster, and some gastric cells cluster with a few BE cells in the ‘enteroendocrine’ cluster. The group described as ‘non-epithelial’ contains some endothelial cells and *CD45*-low immune cells (**Extended Data Figure 4**). Notably, the majority of BE cells (63%) are in the cluster labelled as ‘Barrett’s-type’ that also contains the subset of esophageal cells that have a gene expression profile consistent with their being ESGCs (**Figure 3c,** also see **Supplementary Table 2**). These cells are highly enriched for *LEFTY1*.

**Figure 3.**
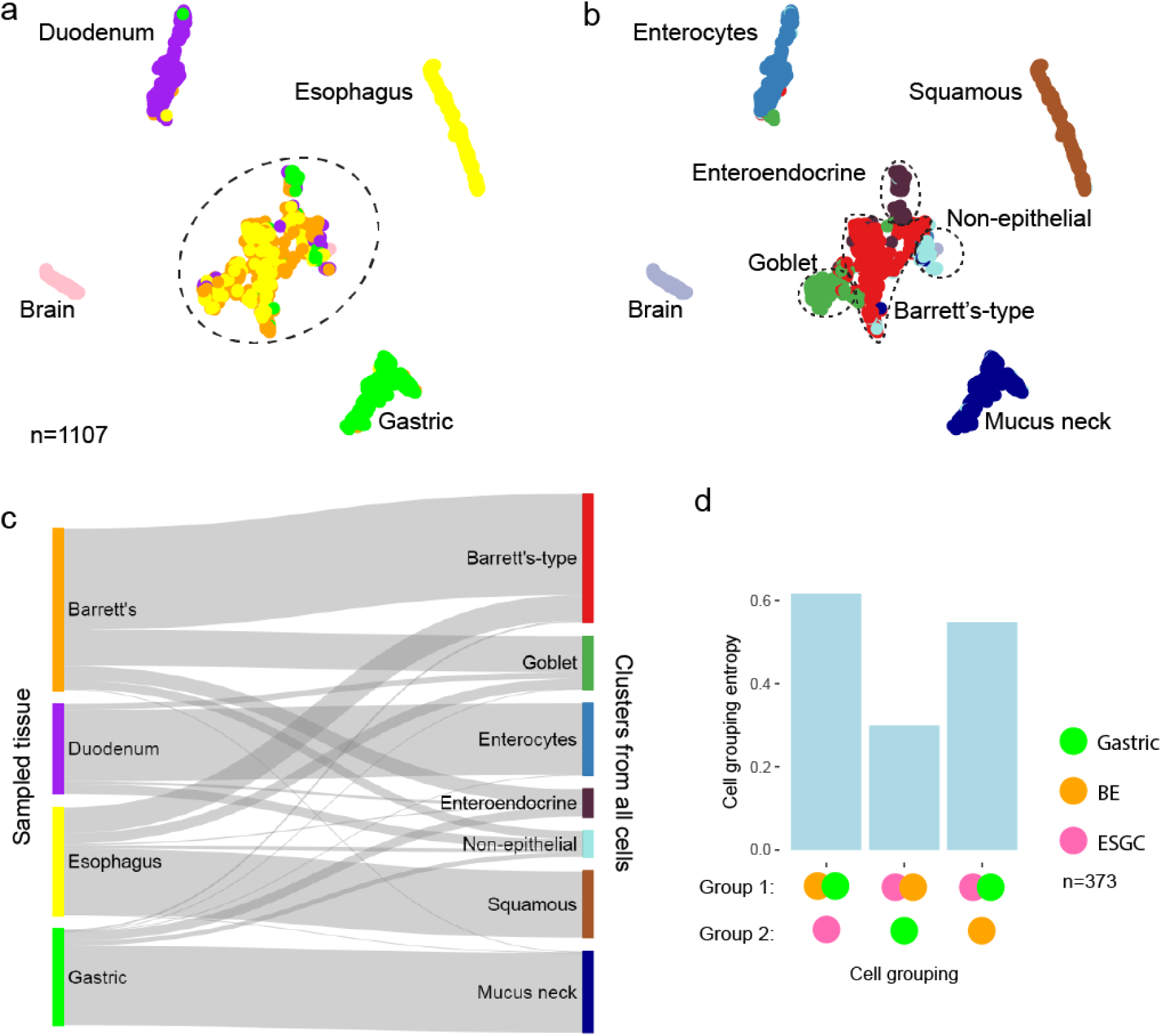
The majority of Barrett’s esophagus cells are transcriptionally similar to esophageal submucosal gland complex (ESGC) cells. **(a)** t-Distributed Stochastic Neighbour Embedding (t-SNE) plots of cells from all samples from four BE patients (n=1107 including brain control), showing similarity of cells in two dimensions, coloured by tissue type (yellow, esophagus; green, gastric cardia; purple, duodenum; orange, Barrett’s esophagus). Dashed circle indicates a group of cells with gene expression consistent with secretory cell function. Brain was used as a control. **(b)** t-SNE plot of cells from all BE patient samples, as in **a**, coloured by how cells contribute to clusters generated by SC3 analysis with 250 repeat clusterings of simulated technical replicates (see Methods, colours as in panel **c** right side). Names given to the clusters are based on expression of known marker genes (see text and Extended Data Figure 2). Dashed lines show the approximate boundaries of named clusters within the secretory cell group. (**c)** Sankey diagram showing how each tissue type sampled contributes to the clusters shown in **b**. Colours and labels on the left indicate sampled tissue (as in **a**); colours and labels on the right indicate cluster (as in **b**). **(d)** Entropy scores for each permutation of ‘gland-like’ cells (n=373), which are a sub-set of gastric (n=175), BE (n=78) and esophagus cells (n=120): excluding gastric and BE cells that expressed *CHGA* or *MUC2* (to exclude enteroendocrine and goblet cells, respectively) and excluding esophageal cells that did not express *TFF3* (to exclude squamous cells). Thresholds were set at the tenth centile of cells in which at least one transcript was detected from each gene.

To test whether this relationship between BE and native esophageal cells with columnar characterization was also seen in patients without BE, we clustered all normal esophagus cells from patients with and without BE groups, using genes differentially expressed between the squamous and Barrett’s type groups in Figure 2. This confirmed that these cells were present in every BE patient sampled, and one patient without BE (**Extended Data Figure 5**).

To confirm whether the relationship between BE cells and ESGCs was stronger than associations with other gland-type cells, we looked across the transcriptional relationships of cells from other tissues, i.e. gastric gland cells and BE cells that did not express *CHGA* or *MUC2* (to exclude enteroendocrine and goblet cells, respectively; see **Methods** for thresholding), and esophageal cells that expressed *TFF2* or *TFF3* (to exclude squamous cells). Comparing entropies of different cell type combinations showed that the BE and ESGC combination had much lower entropy, suggesting this is the strongest relationship (**Figure 3d**). t-SNE, with the inclusion of duodenal cells which expressed the highest levels of *MUC6* (to enrich for duodenal Brunner’s gland-type cells) also showed the strongest strong relationship between BE and ESGC cells (**Extended Data Figure 6**). Collectively, these data show that ESGCs have the greatest transcriptional similarity to BE cells.

### ITLN1 and SPINK4 mark early goblet cells

In this study, 19% of BE cells from all patients were classified as ‘goblet’ cells (**Figure 3c** and **Figure 4a**), which is consistent with the requirement in some countries, such as the US^21^, for goblet cells to be present for the diagnosis of BE. Goblet cells are classically defined by morphological appearance and MUC2 expression. Applying a threshold set at the tenth centile to include 90% of cells in which at least one transcript was detected from each gene (to reduce biological noise), we found that *MUC2* transcriptionally co-expressed with intelectin 1 (*ITLN1*) and Kazal type 4 serine peptidase inhibitor (*SPINK4*) in 61% of goblet cells from duodenum, gastric and BE samples (**Figure 4a,b**). ITLN1 and SPINK4 have been previously shown to mark goblet cells in normal gut and some non-gastrointestinal tissues^22,23^, but we observed some cells in each tissue type that uniquely expressed *MUC2*, *ITLN1* or *SPINK4*. Therefore we hypothesized that their expression pattern might mark stages of goblet cell development *in vivo*. To test this we analysed expression of these proteins by immunostaining 5 human intestinal samples (approximately 500 crypts examined in each sample). ITLN1 and SPINK4 co-staining was consistently present near the crypt base, where undifferentiated cells occur, whereas MUC2 staining was in cells toward the centre and top of the crypts, where terminally differentiated cells are found (**Figure 4c**). This suggests that ITLN1 and SPINK4 mark an earlier stage of goblet cell differentiation than MUC2 in intestine.

**Figure 4.**
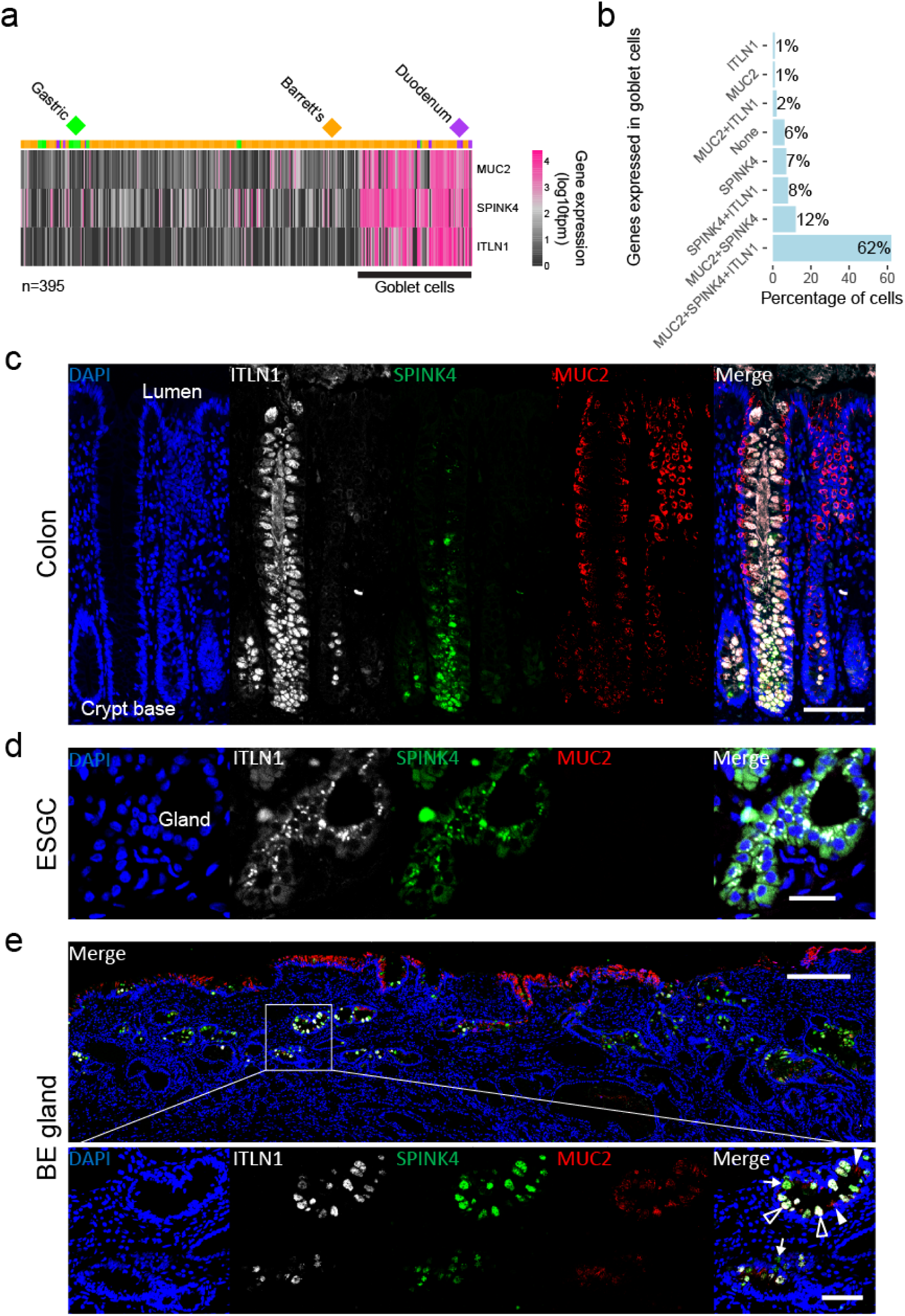
SPINK4 and ITLN1 mark early goblet cells. **(a)** Heatmap of expression of *MUC2*, *SPINK4* and *ITLN1* genes in BE, gastric and duodenal cells in the secretory group (see Figure 2). Tissue of origin of each cell is shown above. The cells in the ‘goblet’ cluster are indicated. **(b)** Bar chart showing the percentage of cells in the ‘goblet’ cell cluster from **a** (n=98) expressing *MUC2*, *ITLN1* or *SPINK4* alone or in different combinations (thresholds set at the tenth centile to include 90% of cells in which at least one transcript was detected from each gene). **(c)** Representative immunofluorescence staining of human colon crypts for MUC2 (red), ITLN1 (white) and SPINK4 (green); nuclei (DAPI) in blue. Scale bar indicates 100 μm. **(d)** Immunofluorescence staining of a representative ESGC beneath BE with MUC2 (red), ITLN1 (white) and SPINK4 (green). Scale bar indicates 20μm. **(e)** Immunofluorescence staining of a representative BE section (scale bar indicates 250 μm). Enlarged images with MUC2 (red), ITLN1 (white) and SPINK4 (green) are shown below (scale bar indicates 50 μm). White tailed arrows show SPINK4 or ITLN1 expressing cells, white filled arrowheads show MUC2 expressing cells, white unfilled arrowheads indicates cells co-staining for MUC2, SPINK4 and ITLN1.

In ESGCs present beneath the mucosa in sections from three patients with BE, we observed that acinar cells consistently co-expressed ITLN1 and SPINK4 without MUC2 (**Figure 4d**). In 30 BE sections from 16 patients we also consistently observed cells expressing ITLN1 or SPINK4 without MUC2 (**Figure 4e,** also see **Extended Data Table 3**). Specifically, in specimens from 5 patients, 41% of MUC2 low cells expressed SPINK4 and/or ITLN1, whereas 28% of cells expressed MUC2 alone (**Extended Data Table 4**). These data suggest that ESGCs and BE may contain early goblet cells, as seen in the colon, and that ITLN1 or SPINK4 might mark cells with some goblet cell characteristics that are not yet morphologically identifiable as goblet cells.

### *OLFM4* marks stem-like transcriptional behaviour in columnar esophageal epithelium

A recent study showed that BE contains pluripotent cells^24^. We therefore analysed all BE and ESGC cells using StemID, which is a published workflow designed to find cells with stem-like properties in single cell RNA-seq data by calculating a ‘stem-ness’ score based on the entropy of cell clusters and the number of links between clusters^25,26^. As a control we analysed duodenum cells and found the highest scoring cluster (**Extended Data Figure 7a,b**, black asterisk) was enriched for *LGR5* expression (**Extended Data Figure 7c,d**), consistent with *LGR5* being a known marker of intestinal stem cells^27,28^. The highest scoring cluster in StemID analysis of BE and ESGC cells together was enriched for expression of the stem-cell associated gene *OLFM4* (**Figure 5a-c**, blue asterisk and **Extended Data Figure 7c**). The second highest scoring cell cluster (**Figure 5a,b,d**, red asterisk and **Extended Data Figure 7c**) was enriched for *LYZ*, a marker of Paneth cells, which are long-lived secretory cells found adjacent to the stem cell niche in the intestinal crypt base. *OLFM4* has been shown to associate with *LGR5* expression and mark stem cells in intestinal tissue in normal and metaplastic contexts^29,30^. Consistent with this, we detected OLFM4 protein in human colon crypt bases, where stem cells are known to be located (**Figure 5e**). In 8 BE sections from 7 patients, we observed that OLFM4 protein expression was less restricted to the crypt base (**Figure 5f, top**), similar to previous observations of LGR5 expression patterns in BE^11^. In ESGCs beneath normal squamous epithelium, OLFM4 positive cells were seen within the gland structures (**Figure 5f, bottom**).

**Figure 5.**
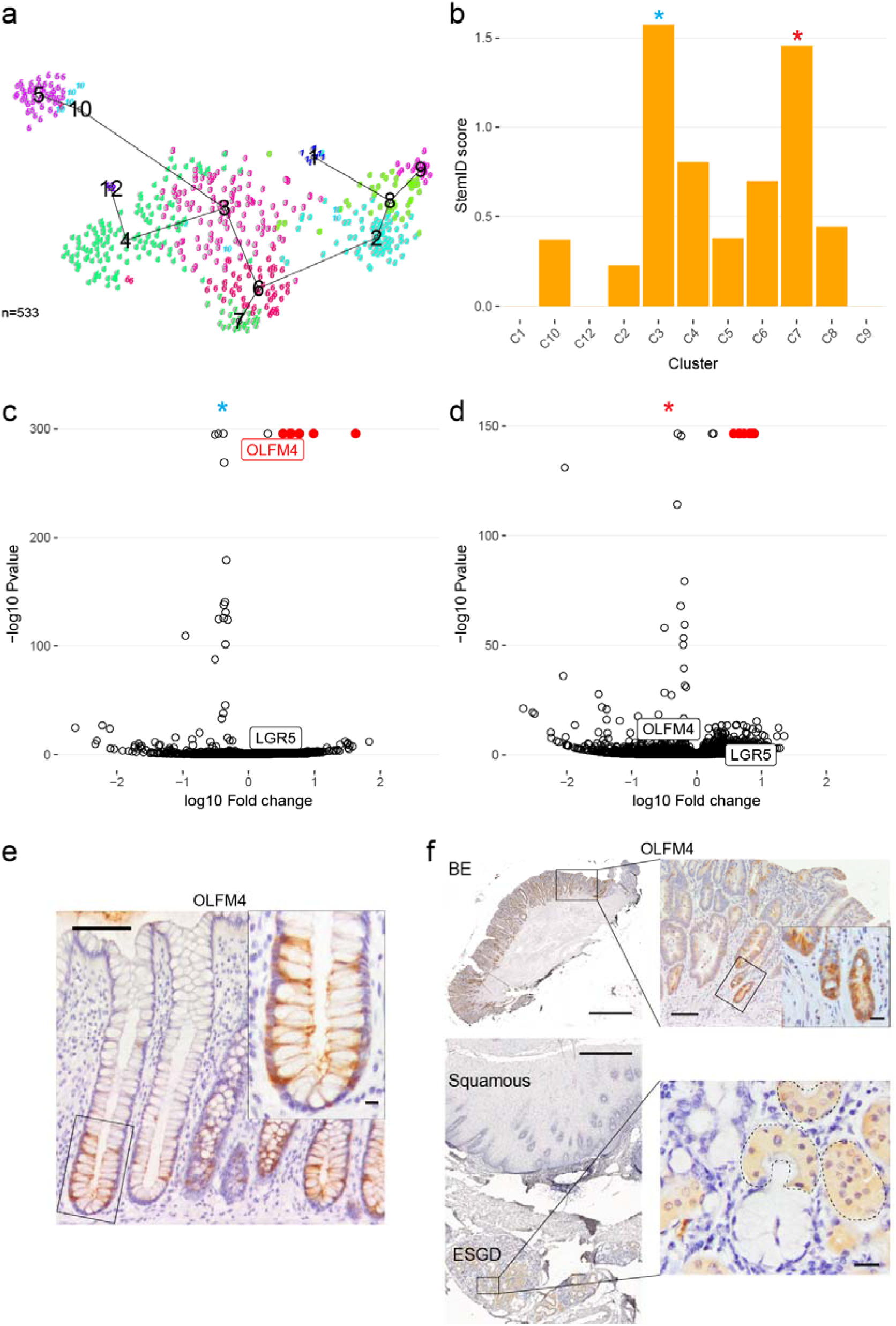
*OLFM4* is upregulated in BE and ESGC cells with stem-lie transcriptional behaviour. **(a)** Plot showing entropy and links between RaceID2 clusters (numbered) as computed by StemID (for more information see: https://github.com/dgrun/StemID) applied to all non-squamous esophageal cells (BE and esophageal cells with <5 KRT14 counts to exclude squamous cells, n=533). **(b)** StemID scores across all clusters computed for the cells in **a**. Scores are calculated from multiplication of the entropy (spread from the cluster mean) and the number of cluster links arising from a given cluster. The highest and second highest scoring clusters are indicated by a blue and red asterisk, respectively. **(c)** Volcano plot of genes from the highest scoring cluster in **b** (C3). Points coloured red indicate the most significant genes with a fold change greater than 2. LGR5 and OLFM4 are labelled in each plot (see Extended Data Figure 7c for more details). **(d)** As in **c**, showing the second highest scoring cluster in **b** (C7). **(e)** Immunohistochemical staining of OLFM4 in human colon (close-up of base of crypt inset). Scale bars indicate 100 μm main image, 10 μm inset. **(f)** Immunohistochemical staining of OLFM4 in BE (top, scale bar indicates 1000 μm), and ESGCs beneath squamous epithelium (bottom, scale bar indicates 500 μm), with enlargements (scale bars indicate 100 μm (top) and 10 μm (bottom), inset of BE gland 20 μm). Black dotted lines (lower right) indicate positively stained areas of OLFM4 in ESGCs.

Notably, *OLFM4* has higher mean expression in the *LEFTY1*-positive clusters (B2/B3) compared to the clusters expressing known markers of the differentiated goblet (*MUC2*) and enteroendocrine (*CHGA*) lineages (**Figure 2a**, B1 and B4, respectively). To examine co-expression of *OLFM4*, *LEFTY1*, *MUC2* and *CHGA* in individual cells we applied a threshold at the tenth centile to include 90% of cells in which at least one transcript was detected from each gene. Using this threshold, half of the BE cells express *LEFTY1* and *OLFM4*, alone or in combination (29% *OLFM4* and *LEFTY1*; 13% *OLFM4* only; 11% *LEFTY1* only). *LEFTY1* and *OLFM4* positive BE cells rarely co-expressed *MUC2* or *CHGA* (**Extended Data Figure 7e)**. Together, these data suggest that B2/B3 represent a cell population that harbours BE progenitor cells.

## Discussion

Our single cell RNA-seq data has resolved cell sub-populations in gastrointestinal epithelia and shown a profound transcriptional similarity between ESGC cells and the largest sub-population of BE cells. This is supported by our observation that this sub-population of BE cells and ESGCs expresses the stem cell-associated gene *OLFM4*, consistent with the notion that these populations might contain similar progenitor cells. Our findings support a potential model in which acid reflux-induced damage to the esophagus is ‘repaired’ by the expansion or selection of ESGCs, which have alkaline secretions and are thus able to play a role in protecting the esophagus from gastroesophageal reflux damage. Further consideration of the functional overlap of other secretory structures with BE and ESGCs, such as salivary and mammary glands may help develop our understanding of the adaptive response to injury that drives metaplasia.

During development of the esophagus, glandular epithelial cells are replaced by squamous epithelium and it has been suggested that the ESGCs can be viewed as a developmental ‘remnant’^31^. This is consistent with our observations of expression of the developmental gene *LEFTY1* in ESGCs and BE. Notably, *LEFTY1* is regulated by TGF-β signalling^32^ and TGF-β is often perturbed in BE^33^, so it will be interesting to explore this relationship further.

Given that rodents lack ESGCs, and the lack of an *in vitro* model of human esophageal glands, analysis of human biopsies currently provides the most reliable approach to dissect the cell relationships of BE. Future improvements in single cell DNA sequencing techniques may enable more systematic genetic confirmation of the cellular origin of BE. Also, it is important to note that our study has not investigated potential origins of EAC. Future studies are needed to address the cell relationships of BE and EAC and how this relates to recent work suggesting that EAC is highly similar to a sub-set of gastric cancers^34^.

We showed that SPINK4 and ITLN1 seem to identify an earlier stage of intestinal metaplasia than marked by MUC2, given that they are expressed lower in intestinal crypts than MUC2 and can be seen without MUC2 in BE. Of clinical importance, our results suggest that intestinal goblet cell characteristics exist even without the presence of morphologically identifiable goblet cells, supporting the view that diagnosis of BE should not require the detection of goblet cells. Together, our findings help characterize BE in humans and will have clinical implications by providing a molecular basis to improve diagnosis of BE. In addition, this study demonstrates the power of single cell analysis of clinical samples to uncover biological relationships among cell types and cellular heterogeneity in healthy and diseased tissues.

---

**Extended Data Figure 1.**
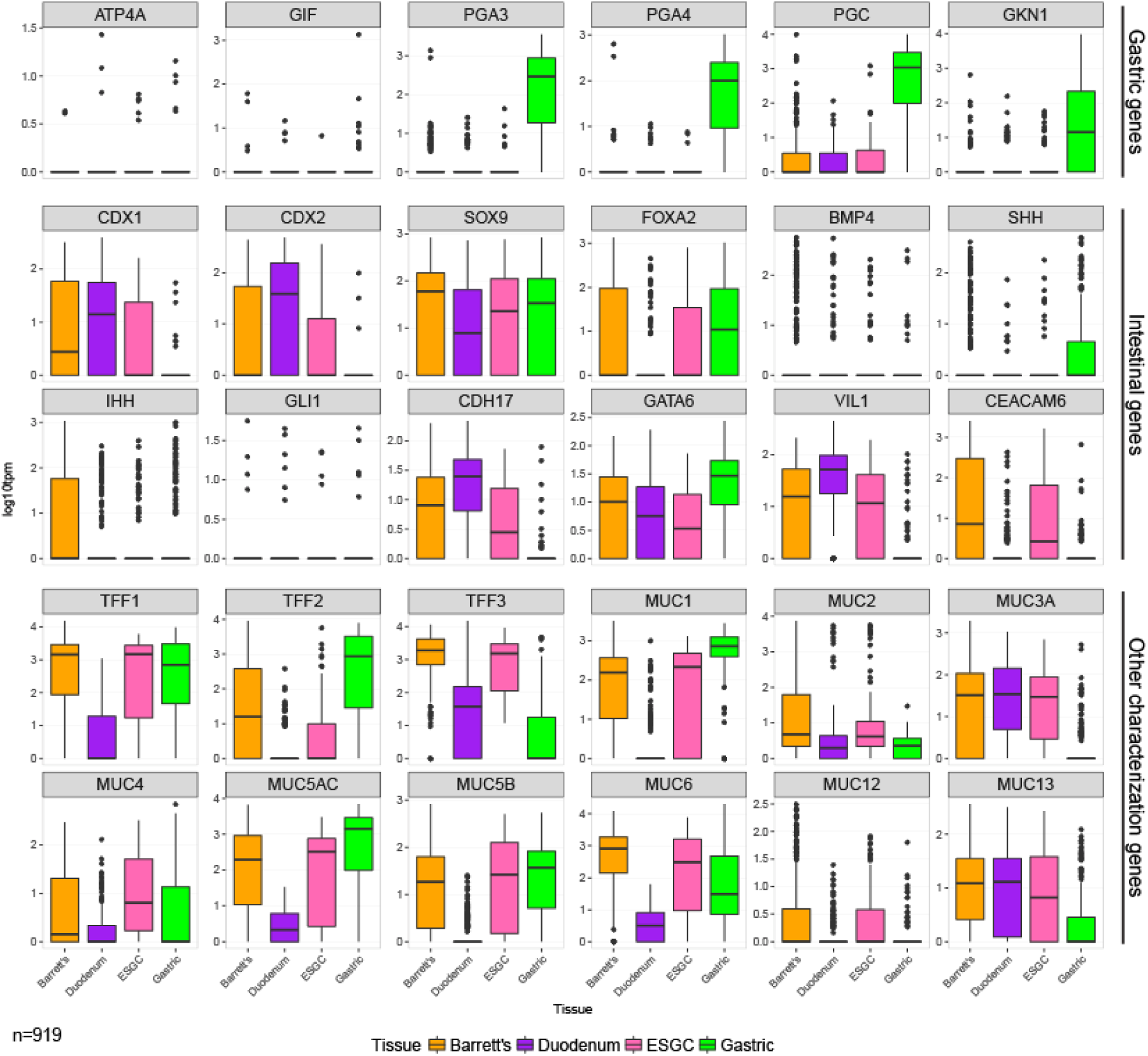
Expression of selected tissue and cell defining genes in columnar cells. Boxplots showing the expression of a selection of genes used to define gastric or intestinal tissue cell types (top two sets of plots labelled gastric genes and intestinal genes on right side), and a panel of mucin and trefoil factor genes (lower set of plots, labelled ‘other characterization genes’). Cells included are from all duodenum (n=207), gastric (n=221) and Barrett’s samples (n=371), columnar type esophageal cells are also included as ‘ESGCs’ (n=120, as in Figure 3). Total n=919.

**Extended Data Figure 2.**
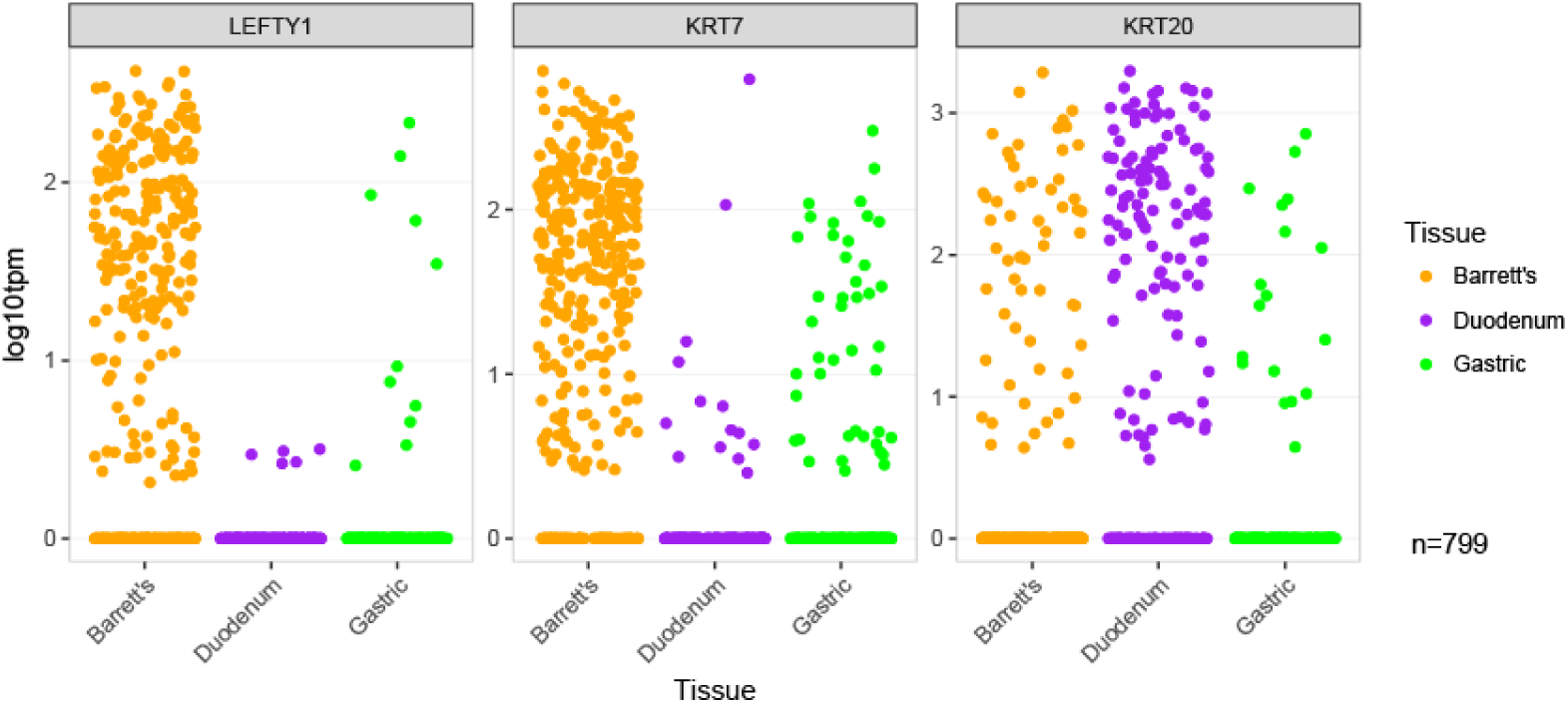
Expression of *LEFTY1, KRT7* and *KRT20* in BE, duodenal and gastric cells. Jitter plots showing the expression of LEFTY1, KRT7 and KRT20 in each included Barrett’s, duodenum and gastric cell.

**Extended Data Figure 3.**
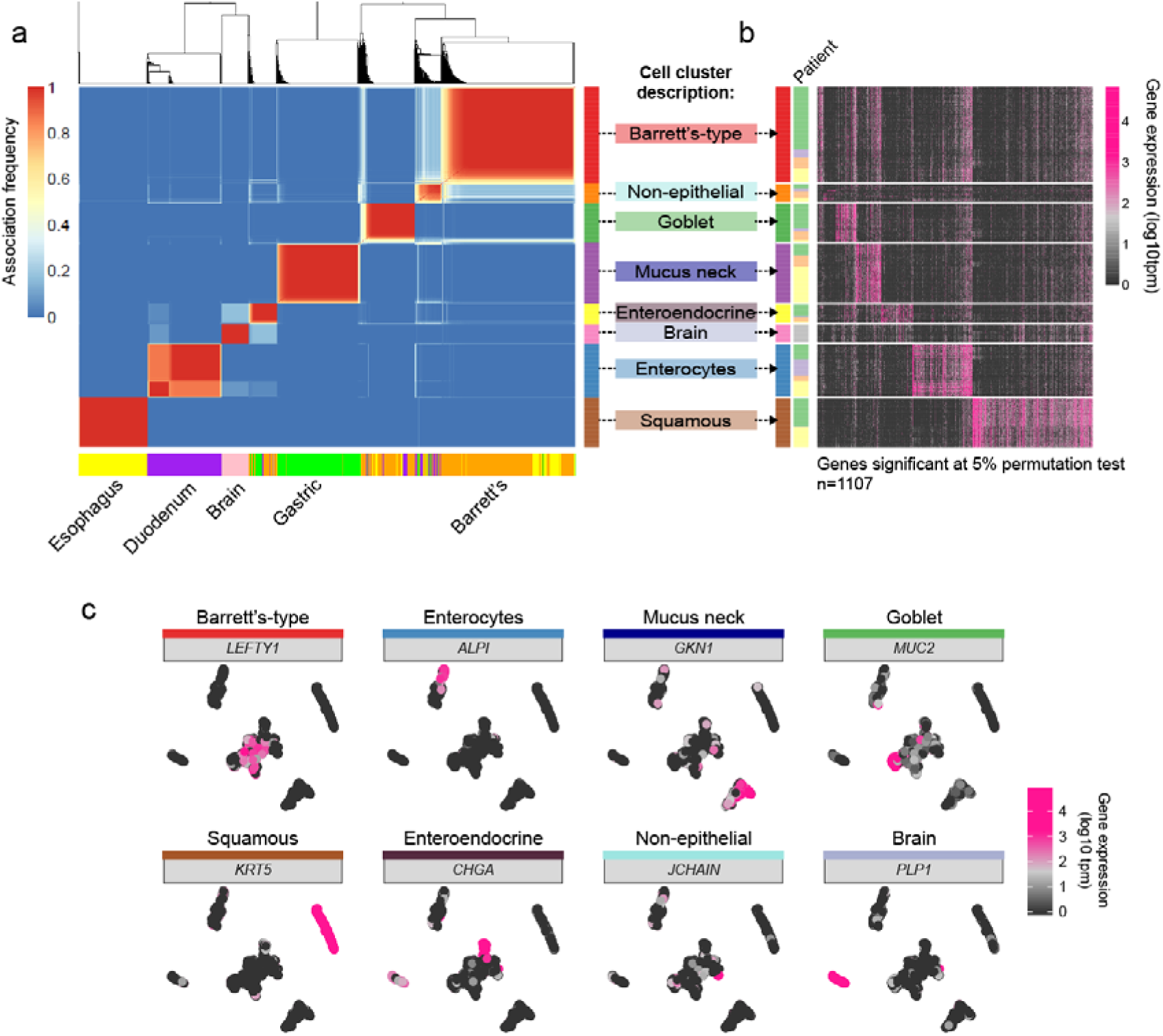
Clustering and differential gene expression profiles of all single cells from Barrett’s esophagus patient samples. **a,** Cluster consensus matrix for single cells from all tissue sites in BE patients (n=1107 including brain positive controls). Blue-to-red colours denote the frequency with which cells are grouped together in 250 repeat clusterings of simulated technical replicates (see Methods). Tissue type is indicated below. Cell clusters are labelled on the right with the cell type they contain or a descriptive term if that cell type has not been previously characterised. **b**, Heatmap showing differentially expressed genes in each cluster from panel **a**, (linked by black dotted arrows). Genes in the heatmap have a minimum of 4 fold higher expression in cells of a given cluster compared to all other cells. Cells from each patient are indicated by coloured bar on the left. **c**, t-SNE plots of all cells from BE patients (n=1107 including positive controls), coloured in each panel by level of expression of a gene that is highly expressed in a particular cluster type (cluster name and gene name shown above plots).

**Extended Data Figure 4.**
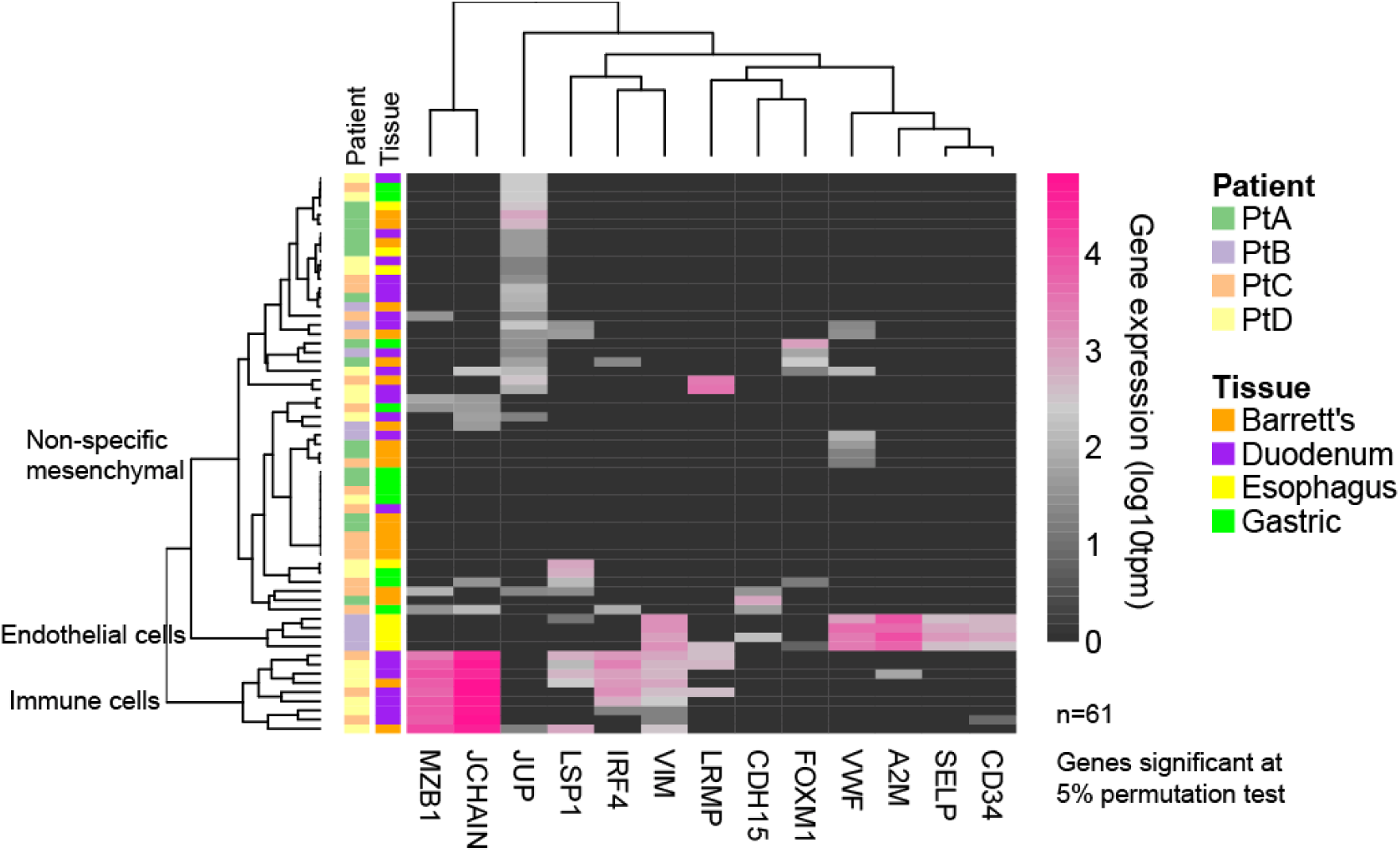
Characterisation of ‘non-epithelial’ cells by gene expression: Heatmap of gene expression in cells in the cluster labelled as ‘non-epithelial’, showing genes significantly upregulated in this cluster (>4 fold, genes significant at 5% permutation test). Genes are clustered by nearest neighbor and the dendrogram is labelled by broad cell-type category based on known functions of the expressed genes. For each cell, tissue type and patient is indicated by the coloured bars on the left.

**Extended Data Figure 5.**
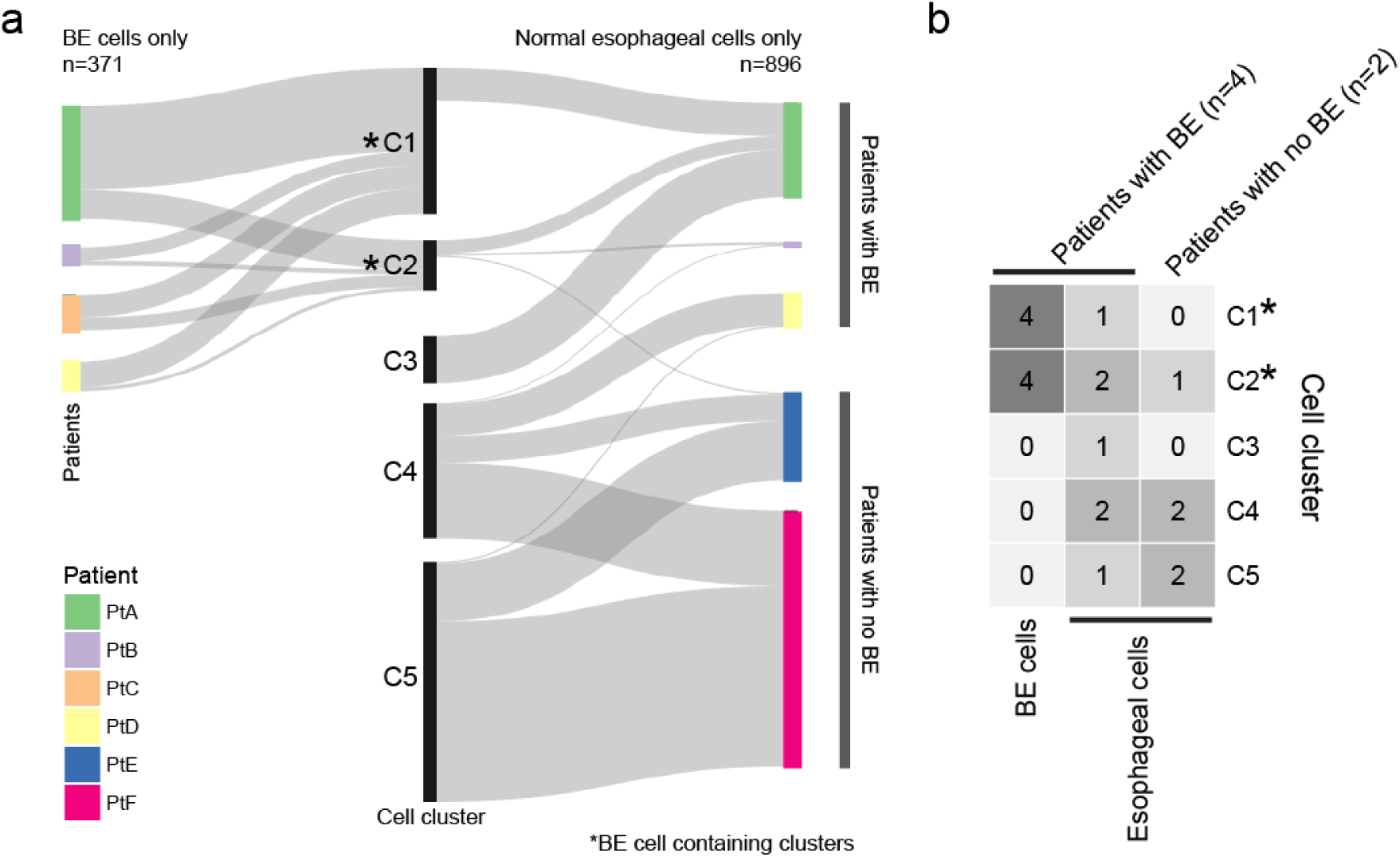
Contribution of cells from different patients to clusters containing Barrett’s esophagus and esophageal cells. **a**, Sankey diagram showing how Barrett’s cells (left) and esophageal cells (right) from each patient contribute to SC3 clusters generated using Barrett’s esophagus and esophageal cells only. Starred clusters (A and B) account for all Barrett’s esophagus cells. **b**, Annotated heatmap showing the number of patients that contribute cells to each of the SC3 clusters (C1- C5, in panel **a**), with contributions separated by tissue origin (BE or esophageal cells) and by patients with BE (n=4) or patients without BE (n=2).

**Extended Data Figure 6.**
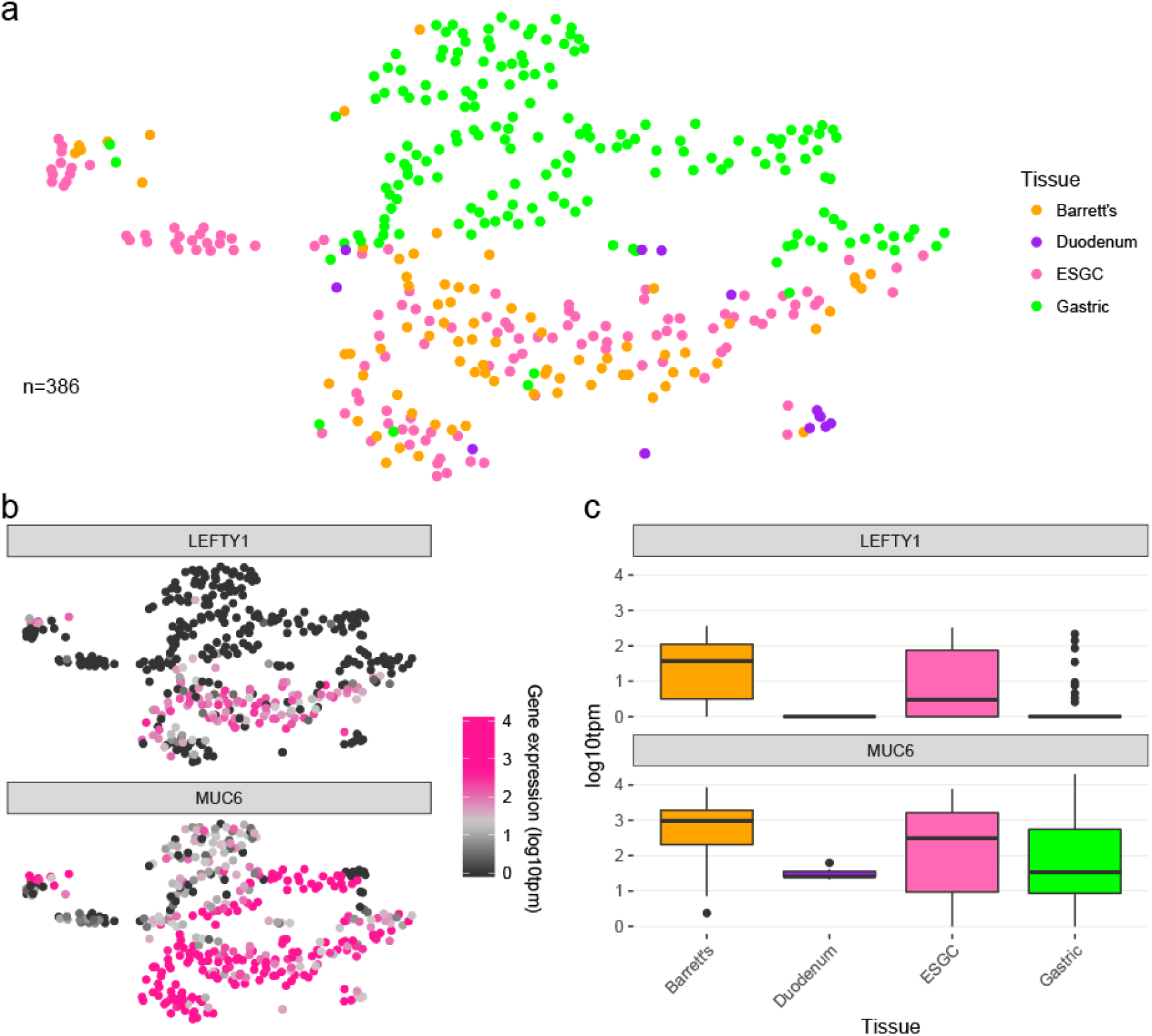
The majority of Barrett’s esophagus cells are transcriptionally similar to esophageal submucosal gland complex (ESGC) cells. **(a)** t-SNE plot of ‘gland-like’ cells (n=386), which are a sub-set of gastric (n=175), BE (n=78), duodenal (n=13) and esophagus cells (n=120): excluding gastric and BE cells that expressed *CHGA* or *MUC2* (to exclude enteroendocrine and goblet cells, respectively), including duodenal cells expressing maximal *MUC6* (to enrich for Brunner’s gland type cells), and excluding esophageal cells that did not express *TFF3* (to exclude squamous cells). Thresholds were set at the tenth centile of cells in which at least one transcript was detected from each gene (with the exception of duodenum cells, in which the threshold was set at the ninetieth centile of *MUC6* expression). **(b)** Expression of *LEFTY1* and *MUC6* in cells shown in **a**. **(c)** Boxplots of expression of LEFTY1 and MUC6 in all columnar cells (see Extended Data Figure 1 for cell selection).

**Extended Data Figure 7.**
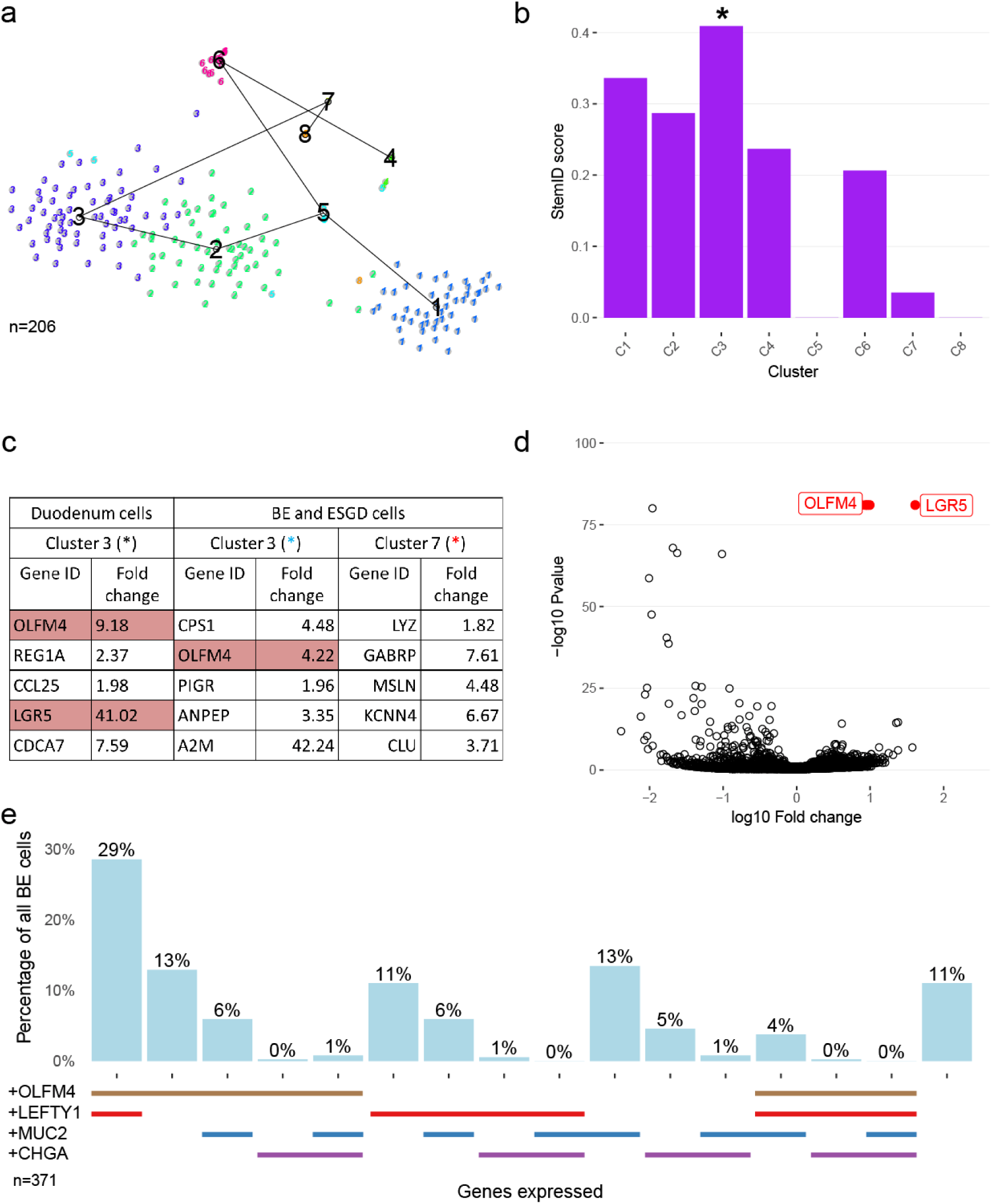
StemID applied to human duodenum, and BE and ESGC cells. **a**, (Top) Plot showing entropy and links between RaceID2 clusters (numbered) as computed by StemID (for more information see: https://github.com/dgrun/StemID) applied to duodenum cells (n=206). **(b)** StemID scores across all clusters computed for duodenum cells. Scores are calculated from multiplication of the entropy (spread from the cluster mean) and the number of cluster links arising from a given cluster. Highest scoring cluster is indicated by a black asterisk. **(c)** A table of the top five most significantly differentially expressed genes in the asterisked clusters (from **b** and Figure 5) with fold change values (against all other clusters) shown. Genes discussed in the text are highlighted. **(d)** Volcano plot of genes from the highest scoring cluster from duodenum cells (C3). Points coloured red indicate the most significant genes with a fold change greater than 2. Selected genes of interest are labelled. **(e)** Bar chart showing percentage of cells expressing *OLFM4*, *LEFTY1*, *MUC2* or *CHGA* alone and in all combinations (thresholds for calling a gene ‘expressed’ were set at the tenth centile to include 90% of cells in which at least one transcript was detected from each gene).

**Extended Data Figure 8.**
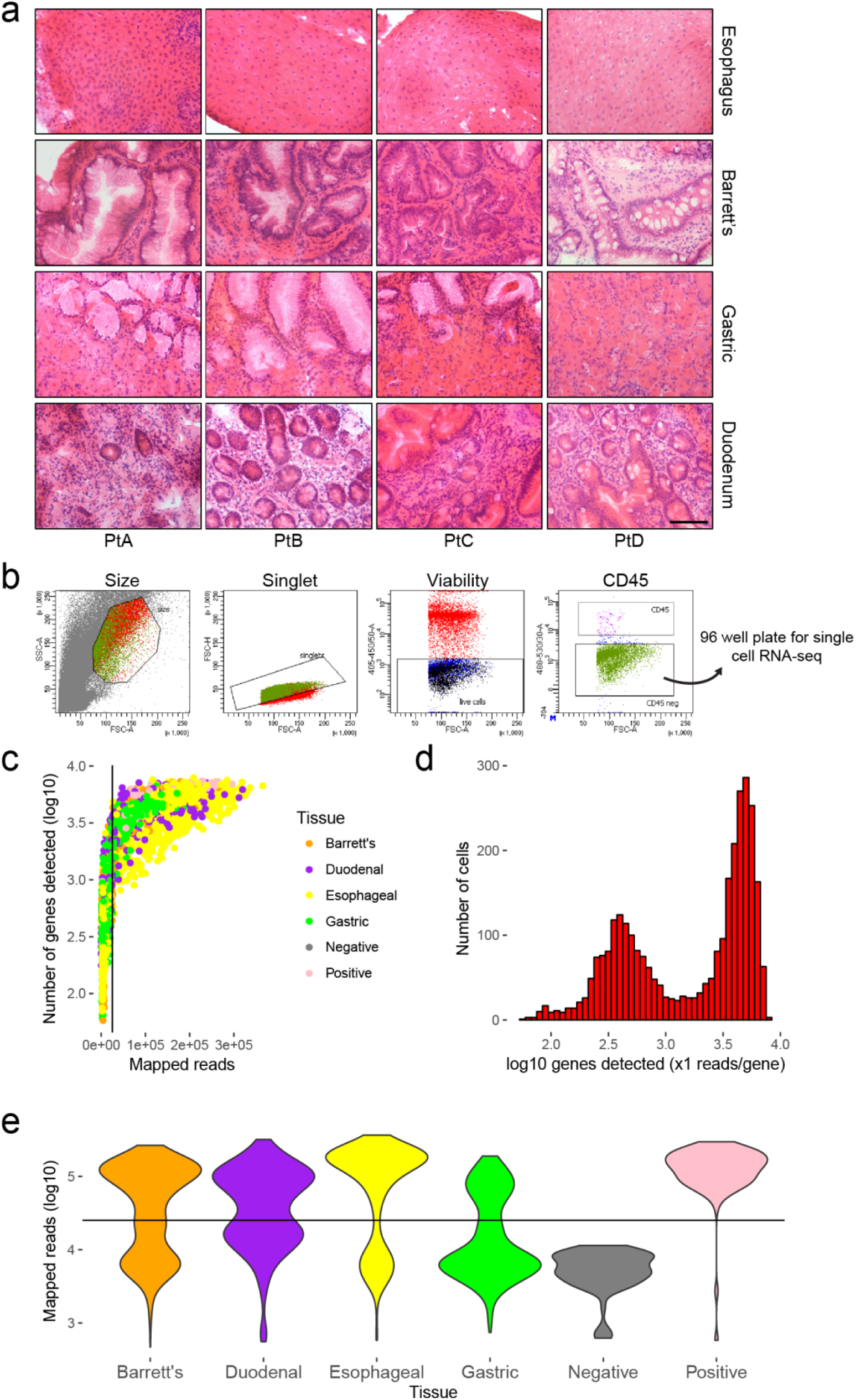
Sample analysis and quality control. **a**, Haematoxylin and eosin staining of frozen sections taken from the biopsy fragments that were used for single cell and bulk RNA-sequencing from four patients (excluding the normal esophageal samples taken from non-Barrett’s esophagus patients) (Scale bar represents 100 μm). **b**, An example of a FACS sort for a duodenal sample. Cells were selected on size gating, singlet gating, viability (DAPI negative), and CD45 negative gating to exclude leukocytes. **c**, Plot of number of genes detected (at least one read per gene) against total mapped reads for each cell (n=3051). Each dot indicates a cell. Black line indicates the mapped read threshold determined by the bimodality of positive control (brain RNA diluted to 10pg) and negative control (lysis buffer only in well) read mapping. **d**, Histogram of the distribution of number of detected genes across all cells. **e**, Distribution of total mapped reads per cell, separated by tissue. Horizontal black line indicates the mapped read threshold as in **c**.

**Extended Data Table 1.** Sampled patients. BMI, body mass index; PPI, protein pump inhibitor treatment. Prague indicates Prague classification for measuring the length of Barrett’s esophagus, where C is proximal extent of circumferential lesion and M is maximal proximal extent (na for patients without BE). scRNA-seq and Bulk RNA-seq indicate whether samples from each patient were used for single cell or bulk RNA-seq, respectively.

| Patient | Age | Sex | Smoking (cigs/day) | Alcohol (units/week) | BMI | PPI | Prague | scRNA-seq | Bulk RNA-seq |
| --- | --- | --- | --- | --- | --- | --- | --- | --- | --- |
| PtA | 78 | M | 0 | 21 | 26.2 | Yes | C3M5 | Yes | Yes |
| PtB | 77 | M | 0 | 4 | 25.7 | Yes | C0M3 | Yes | Yes |
| PtC | 47 | F | 0 | 16 | 22.2 | Yes | C0M2 | Yes | Yes |
| PtD | 90 | F | 0 | 6 | 23.7 | Yes | C9M11 | Yes | Yes |
| PtE | 82 | F | 0 | 0 | 27.1 | Yes | na | Yes | No |
| PtF | 68 | F | 0 | 0 | 25.8 | Yes | na | Yes | No |
| PtG | 69 | F | 0 | 0 | 32.6 | Yes | C5M7 | No | Yes |
| PtH | 86 | M | ex 10/day | 35 | 21.9 | Yes | C8M10 | No | Yes |
| PtI | 74 | F | 0 | 0 | 35.7 | Yes | C1M3 | No | Yes |

Prague score: C = Circumferential extent of lesion; M = Maximal extent of lesion

**Extended Data Table 2.** Summary of immunohistochemical staining of BE specimens. Integers denote the number of sections (one section per slide) immunohistochemically stained for each antibody. A total of 81 sections were stained from 19 patients. Basic pathological details are noted (BE, Barrett’s esophagus; IM, intestinal metaplasia; LGD, low grade dysplasia; HGD, high grade dysplasia).

| Patient sample # | Histological diagnosis | Number of sections stained for each antibody |  |  |  |  |  |  |
| --- | --- | --- | --- | --- | --- | --- | --- | --- |
|  |  | OLFM4 | LEFTY | KRT7 | CHGA | MUC2 | SPINK4 | ITLN1 |
| 7825 | BE with IM | 1 | 1 | 1 | 1 | 1 | 0 | 0 |
| 5368 | BE with IM | 1 | 1 | 1 | 1 | 1 | 0 | 0 |
| 3104 | BE with IM some LGD | 2 | 1 | 1 | 1 | 1 | 0 | 0 |
| 3612 | BE with IM | 1 | 1 | 1 | 1 | 1 | 0 | 0 |
| 813 | BE with IM some LGD | 1 | 0 | 0 | 1 | 0 | 2 | 2 |
| 8226 | BE with IM | 1 | 0 | 0 | 2 | 0 | 2 | 2 |
| 2318 | BE with IM | 1 | 0 | 0 | 1 | 0 | 1 | 1 |
| 8173 | BE with IM some LGD | 0 | 1 | 0 | 1 | 0 | 3 | 3 |
| 7974 | BE with IM | 0 | 2 | 0 | 1 | 1 | 1 | 1 |
| 282 | BE with IM | 0 | 0 | 0 | 2 | 1 | 1 | 1 |
| 774 | BE with IM | 0 | 0 | 0 | 1 | 1 | 1 | 1 |
| 5519 | BE with IM | 0 | 0 | 0 | 1 | 1 | 1 | 1 |
| 5097 | BE with IM dysplastic foci | 0 | 0 | 0 | 2 | 0 | 2 | 2 |
| 6446 | BE with IM some LGD | 0 | 2 | 0 | 0 | 0 | 0 | 0 |
| 5073 | BE with IM | 0 | 2 | 0 | 0 | 0 | 0 | 0 |
| 9017 | BE with IM | 0 | 2 | 0 | 0 | 0 | 0 | 0 |
| 7836 | BE with IM some LGD and HGD | 0 | 2 | 0 | 0 | 0 | 0 | 0 |
| 6198 | BE with IM some LGD | 0 | 1 | 0 | 0 | 0 | 0 | 0 |
| 6114 | BE with some LGD | 0 | 0 | 0 | 1 | 0 | 0 | 0 |

**Extended Data Table 3.** Summary of immunofluorescent staining of BE specimens. Integers denote the number of sections (one section per slide) triple stained for ITLN1, MUC2 and SPINK4. A total of 30 sections were stained from 16 patients. Asterisked patient sample #s indicate samples which were also used for immunohistochemical staining (see Extended Data Table 3). Basic pathological details are noted (BE, Barrett’s esophagus; IM, intestinal metaplasia; LGD, low grade dysplasia; HGD, high grade dysplasia).

| Patient sample # | Histological diagnosis | Number of sections stained for ITLN1/SPINK4/MUC2 |
| --- | --- | --- |
| 8173 | BE with IM | 3 |
| 8226 | BE with IM | 3 |
| 5368 | BE with IMsome LGD | 3 |
| 6198 | BE with IMsome LGD | 3 |
| 8720* | BE with IM | 3 |
| 452* | BE with IM | 3 |
| 3458* | BE with IM | 2 |
| 9811* | BE with IM | 2 |
| 5097 | BE with IMsome LGD | 1 |
| 6773* | BE with IMsome LGD | 1 |
| 3612 | BE with IMsome LGD | 1 |
| 7974 | BE with IM | 1 |
| 6446 | BE with IMsome LGD | 1 |
| 813 | BE with IMsome LGD | 1 |
| 2318 | BE with IM | 1 |
| 7709* | BE with IMsome LGD and HGD | 1 |

**Extended Data Table 4.** MUC2, ITLN1 and SPINK4 expression from five BE patients. n is the number of cells scored as positive for expression of each protein by immunofluorescence. % is the percentage of the total number of scored cells (bottom row) that is positive for the protein(s) listed. See Methods for details of the antibodies used.

|  |  |  | Patient sample # |  |  |  |  |  |  |  |  |  |
| --- | --- | --- | --- | --- | --- | --- | --- | --- | --- | --- | --- | --- |
| Antibody |  |  | 8173 |  | 8226 |  | 5097 |  | 6446 |  | 5073 |  |
| MUC2 | SPINK4 | ITLN1 | n | % | n | % | n | % | n | % | n | % |
| High | High | High | 22 | 6 | 34 | 12 | 41 | 16 | 15 | 5 | 2 | 3 |
| High | High | Low | 28 | 7 | 26 | 9 | 6 | 2 | 190 | 59 | 4 | 5 |
| High | Low | High | 11 | 3 | 17 | 6 | 4 | 2 | 2 | 1 | 3 | 4 |
| High | Low | Low | 150 | 38 | 54 | 18 | 30 | 12 | 55 | 17 | 44 | 55 |
| Low | High | High | 40 | 10 | 134 | 45 | 70 | 28 | 26 | 8 | 11 | 14 |
| Low | High | Low | 110 | 28 | 11 | 4 | 7 | 3 | 20 | 6 | 4 | 5 |
| Low | Low | High | 16 | 4 | 7 | 2 | 90 | 36 | 11 | 3 | 7 | 9 |
| Low | Low | Low | 17 | 4 | 12 | 4 | 5 | 2 | 5 | 2 | 5 | 6 |
| Total cells |  |  | 394 |  | 295 |  | 253 |  | 324 |  | 80 |  |
| MUC2 Low, SPINK4 and/or ITLN1 High |  |  | 166 | 42 | 152 | 52 | 167 | 66 | 57 | 18 | 22 | 28 |
| MUC2 High, SPINK4 and ITLN1 Low |  |  | 150 | 38 | 54 | 18 | 30 | 12 | 55 | 17 | 44 | 55 |

**Supplementary Table 1. Differentially expressed genes between tissue specific cell clusters.**

Tables containing differentially expressed genes between clusters for each of Barrett’s oesophagus, duodenum, gastric and oesophagus cells in patients with BE. Differentially expressed genes were obtained by comparing cells of each cluster against all other clusters in a given tissue type. See **Methods** for details on how clusters were determined, and thresholds for significance were determined. Results from ANOVA–type analysis have been excluded from this table.

**Supplementary Table 2. Differentially expressed genes between all tissue clusters.**

Tables containing differentially expressed genes between clusters for all cells analysed in patients with BE. Differentially expressed genes were obtained by comparing cells of each cluster against all other clusters. See **Methods** for details on how clusters were determined, and thresholds for significance were determined. Results from ANOVA–type analysis have been excluded from this table.

## Methods

### Sampling

Patients attending routine endoscopic surveillance of BE and patients with mild reflux symptoms undergoing gastroscopy for diagnostic purposes gave informed consent and provided samples (South Central - Oxford C Research Ethics Committee: 09/H0606/5+5). Patient numbers were chosen to provide suitable biological replicates, and cells sequenced to provide balanced sample sizes at sequencing input. Double bite quadrantic 2mm biopsies were obtained endoscopically using standard biopsy forceps (Radial Jaw 4 Standard Capacity, Boston Scientific, Natick, USA) from a central region of the BE segment avoiding the proximal BE margin as well as the oesophagogastric junction. Control samples were taken from the second part of the duodenum, the stomach 20mm distal to the gastroesophageal junction and the normal esophageal squamous epithelium at least 20mm clear of the most proximal extent of BE. Each sample was fragmented and then pooled to ensure all sampling sites were represented in each investigative modality. Fragments pools were divided into three groups for histological verification, whole-tissue RNA-seq and single cell RNA-seq (**Figure S6a**). Patients were selected based on their previously known pathological features (**Table S1**).

### Cell isolation

Sample fragments were placed directly into digestion solution, made with 1x phosphate buffered solution (Gibco™), heat inactivated fetal bovine serum (Sigma-Aldrich®), EDTA and type I collagenase (Worthington Biochemical Company®), and gently oscillated for 60 minutes. Samples were then further fragmented with scissors and briefly manually triturated with a p1000. Fragments were allowed to settle and the cell containing supernatant filtered (Sysmex Celltrics® 100 micron) into a 15ml Falcon tube. This process was repeated 3 times and the product centrifuged to create a cell pellet which was resuspended in sorting buffer. A small amount of each sample was pooled for labelling controls. Pre-conjugated CD45-FITC (1:10, mouse monoclonal, cat. 130-080-202, Miltenyi Biotec)^35^ and EpCAM-PE (1:10, mouse monoclonal, cat. 130-110-999, Miltenyi Biotec)^36^ antibodies were added to cell suspensions to identify epithelial and immune cells, respectively, and they were incubated/washed according to manufacturer’s advice. DAPI (1:2000, Sigma-Aldrich®) was added to cell suspensions immediately prior to sort. FACS was carried out using a BD Biosciences FACS Aria IIIu platform with 70μm nozzle. Cells were selected based on size and singlet gating to saturate cell output while minimising debris passed to subsequent gates. High EpCAM+ cells were then selected (where an abundance of cells were available) to ensure epithelial provenance was maximised (**Figure S6b**). Resultant cells were sorted directly into 96 well plates (Life Technologies™ MicroAmp® Optical 96-well Reaction Plate) pre-prepared with 2μl 0.2% Triton™ X-100 (Sigma-Aldrich®) and RNAse inhibitor (Takara Recombinant RNase Inhibitor) at 19:1 and then immediately frozen on dry ice. To confirm spectral accuracy, compensation bead controls and pooled cell suspensions were used for fluorescence-minus-one controls. Each plate was re-permuted to avoid batch effects at the next stages of preparation, with no single plate containing cells from only a single patient or tissue type. Variable patterns of 6 blank wells were also prepared in each plate, 3 of which had a 10pg of brain total RNA (Agilent Technologies) added as a positive control.

### Single cell RNA-seq

Transcriptome libraries were prepared using a Biomek FX liquid handling instrument (Beckman Coulter) with a custom adaptation of the published smart-seq2 method^37,38^, with minor modifications, and Nextera XT (Illumina®) methodology with custom, unique index primers after tagmentation and ERCC spike-in. Libraries were sequenced using the Illumina® HiSeq 4000 platform, aiming for 3.5×10^5^ reads per cell at 75bp paired end.

### Bulk RNA-seq

Tissue fragments were processed using the *mir*VanA^TM^ miRNA Isolation Kit (ThermoFisher) according to manufacturer’s guidance. Total RNA was enriched using ribodepletion (Ribo-Zero, Illumina®) prior to cDNA conversion. Second strand DNA synthesis incorporated dUTP. cDNA was end repaired, A-tailed and adaptor ligated. Samples then underwent uridine digestion. The prepared libraries were size selected and multiplexed before 75bp paired end sequencing using the Illumina® HiSeq 4000 platform.

### Data analysis

All data were mapped using STAR^39^ (release 2.5.2a) to the hg19 version of the human genome with transcriptome annotations from Gencode (release 25). Counts tables were made with HTSeq^40^. Cells were excluded with fewer than 25119 fragments mapping to the transcriptome (a threshold which excludes all negative controls, and includes all positive controls, see **Figure S6c-e**). Counts were TMM-normalised and FPKM values were calculated. Genes with less than 4 FPKM in at least 3 cells were filtered out. After re-normalisation, expression values were converted to TPM. A further gene filtering step was included to remove highly expressed genes with low variability across all samples (cells in the top decile for mean expression and below the fifth centile for coefficient of variation). SC3^41^ was used to provide cell cluster information. Cluster robustness to experimental technical variation was tested using BEARscc^42^ which models technical noise from ERCC measurements. Cluster number, k, was chosen manually using the distribution of cluster-wise mean silhouette widths across clusters in all 250 simulated technical replicates for each cluster number k (2 to 8 for individual tissue and 1 to 15 for all tissues). t-SNE data were generated using the Barnes-Hut implementation of t-SNE^43^ in R. Cluster entropies were calculated using NMF^44^ in R. Differential expression analysis was carried out between cell groups using edgeR^45^ from normalized counts according to the package manual. P values used were determined by permutation test at 5% (250-1000 permutations) to allow for multiple comparisons or, in cases of unbalanced sample numbers, converted to false discovery rates (FDR) by the Benjamini-Hochberg procedure. Identification of stem-like cells was performed using RaceID2 and StemID according to the authors’ recommendations^25,26^. Where gene expression is described in binary terms, the threshold was set to include or exclude 90% of cells with the highest expression of a given gene, to allow for biological noise.

### Immunohistochemistry and immunofluorescence staining of human tissue

Esophageal samples from esophagectomy specimens (5 patients) containing normal mucosa and gland structures and endoscopic mucosal resection specimens (30 patients) with Barrett’s esophagus were obtained from the Oxford biobank. Sections were de-waxed, rehydrated and incubated with 3% hydrogen peroxide in methanol to block endogenous peroxidase activity (10 minutes). Antigen retrieval carried out using pH6 sodium citrate Sections were then blocked with normal goat serum and incubated overnight at 4 °C with a primary antibody against anti-KRT14 (IHC, 1:1000, rabbit polyclonal, cat. PRB-155P, BioLegend), anti-TFF3 (IHC, 1:1000, mouse monoclonal, cat. WH0007033M1, Sigma-Aldrich®)^46^, anti-MUC2 (IHC, 1:300, rabbit polyclonal, cat. SC-15334, Santa Cruz Biotechnology)^47^, anti-CHGA (IHC, 1:500, rabbit polyclonal, cat. ab15160, Abcam)^48^, anti-KRT7 (IHC, 1:4000, rabbit monoclonal, cat. ab181598, Abcam)^49^, anti-LEFTY (IHC, 1:1000, rabbit polyclonal, cat. ab22569, Abcam)^50^, anti-OLFM4 (IHC, 1:200, rabbit monoclonal, cat. D1E4M, Cell Signalling Technology®), anti-ITLN1 (IHC/IF, 1:500, sheep polyclonal, cat. AF4254, R&D systems)^51^, anti-MUC2 (IF, 1:300, mouse monoclonal, cat. ab11197, Abcam)^52^, anti-SPINK4 (IF, 1:500, rabbit polyclonal, cat. HPA007286, Sigma-Aldrich®)^53^. For immunohistochemical staining samples were then treated with biotinylated secondary antibody (Vector Labs; 1:250) for 40 minutes at room temperature. The staining reaction was worked up using the Vector Elite ABC kit and counterstained with haematoxylin. Samples were examined by a pathologist using a histology microscope. For immunofluorescent staining expression was detected using Alexa Fluor (1:250, Molecular Probes) for one hour. DAPI (1:2000, Sigma-Aldrich®) was used to stain nucleic acids. Samples were observed using a confocal microscope system (LSM 710; Carl Zeiss).

## Data availability

Raw data will be available in the database of Genotypes and Phenotypes, following the necessary consents to protect donor anonymity

## Author contributions

R.P.O. and M.J.W. collected biopsy samples and prepared them for sequencing. M.J.W. carried out the immunoreactive staining and imaging. R.P.O. and D.S.T. carried out RNA-seq mapping and data analysis. B.B., A.B., M.M. and N.D.M. helped to design and curate the clinical data and sample collection. R.G. and L.M.W. provided pathological interpretation of all samples used. A.G., P.P. and D.B. generated all sequencing data used. C.P.P. provided computational oversight of the data analysis. B.S.-B. provided overall supervision of the computational analysis of the data and X.L. provided overall supervision of the project. The manuscript was written by R.P.O. and X.L., with assistance from M.J.W., B.S.-B. and D.S.T. Figures were prepared by R.P.O., M.J.W. and D.S.T.

The authors declare no competing financial interests. The views expressed are those of the authors and not necessarily those of the NHS, the NIHR or the Department of Health.

## Acknowledgments

We thank Mary Muers for helping to prepare the manuscript, Andrew Roth for comments on statistical analysis of the data, Sally-Ann Clark and Paul Sopp for providing FACS expertise, John Findlay for help with ethical approval, and Rory Bowden, Amy Trebes and the High-Throughput Genomics team at the Wellcome Trust Centre for Human Genetics, Oxford for assistance with sequencing.

Funding support was from Ludwig Institute for Cancer Research, Cancer Research UK, the Medical Research Council, NIHR Biomedical Research Centre, based at Oxford University Hospitals Trust, Oxford Health Services Research Committee and Oxford University Clinical Academic Graduate School.

